# Supplementing floral resources does not mitigate the negative effects of urbanisation on cavity-nesting bees and wasps

**DOI:** 10.1101/2024.12.19.629413

**Authors:** Emilie E. Ellis, Stuart A. Campbell, Jill L. Edmondson

## Abstract

1. Cavity-nesting bees and wasps provide essential ecosystem services such as pollination and pest control. Within urban environments, it is likely that cavity-nesting insects are using greenspaces to forage for food and nesting resources, and thus may benefit from urban greenspace management interventions, such as the addition of flower patches and “bee hotels” (trap nests).
2. In contrast to the relatively large body of work demonstrating the effects of wildflower provisioning on bees in an agricultural context, there have been few studies testing the benefits of floral provisioning, or the relative importance of nest site and food constraints on urban cavity-nesting bees and wasps. Consequently, we lack an empirical evidence base for optimising and informing conservation interventions for these taxa.
3. To understand how the addition of floral resources influences cavity-nesting insect communities, we conducted a 2-year study in urban greenspaces used for horticulture (allotments). We manipulated floral and nesting resources by adding nectar-rich flower patches and artificial trap nests, and compared nest colonisation by cavity-nesting bees and wasps in sites with floral additions and where no additions were made. Our sites were configured along an urbanisation gradient, allowing us to examine how surrounding urbanisation influenced these patterns.
4. Contrary to our predictions, we found no significant difference in trap nest uptake in sites where floral resources were added, compared to control sites. However, cavity-nesting insect abundance showed a 6-fold decrease over our urbanisation gradient. The addition of floral resources did not mitigate the negative effects of urbanisation on cavity-nesting insects.

**Synthesis and Applications:** Although enhancing floral resources in urban systems is a recognised strategy for conserving key pollinators such as social bees, our findings suggest that this approach alone does not address the challenges faced by cavity-nesting bees or wasps in urban landscapes. Our results suggest that accounting for the diversity of resource requirements and incorporating landscape-scale analysis is essential to ensure local conservation interventions will benefit a diversity of insect communities.

## Introduction

The conservation of insects is vital to maintain ecosystem function and support the delivery of ecosystem services to society. Cavity-nesting hymenopterans (bee and wasp species that nest in cavities above ground) are a diverse group of insects that carry out essential ecosystem functions that are particularly relevant to the effective functioning of human-dependant systems, particularly through pollination and pest control. Cavity-nesting bees provide pollination services to wild and crop species of plants (Winfree et al., 2007; Brittain et al., 2013; Garibaldi et al., 2013, MacIvor et al., 2014) and cavity-nesting wasps are effective predators of arthropods such as lepidopteran caterpillars (Tscharntke et al., 2005; Bianchi et al., 2006; Careless et al., 2014). Adult cavity-nesting wasps, like cavity-nesting bees, also feed on nectar and have been shown to provide an overlooked but effective pollination service (Mello et al., 2011; Brock et al., 2021). Current conservation efforts that enhance the availability of floral resources or provide nesting sites could therefore be an important intervention to maintain or enhance both insect communities. However, the effectiveness of these conservation interventions remains poorly understood for cavity-nesting bees, and virtually unknown for cavity-nesting wasps (Brock et al., 2021).

Urban areas pose a complex set of environmental challenges to pollinating insect communities, and mitigating these challenges has become a focus of both research, and pollinator conservation efforts, including initiatives like “No Mow May,” “Save Our Bees” by the Wildlife Trust, and the U.K. National Pollinator Strategy 2021-2024 (DEFRA, 2022). The process of urbanisation is one of the main drivers of global declines in insect diversity and is largely driven by the conversion semi-natural habitats into cities and towns (Wagner et al 2021). However, greenspaces (e.g. parks, allotments, gardens, woodlands) are an integral part of the fabric of urban ecosystems; for example, in the U.K. they cover approximately 50% of a city’s land area (Edmondson et al., 2020; Dennis et al., 2018). The complex mosaic of greenspaces interspersed across the urban landscape act as refuges for numerous insect taxa (Hall et al., 2012, Baldock et al., 2015; Theodorou et al., 2020). From a conservation ecology perspective these urban greenspaces have become the focus of research due to the active conversion of homogenous areas (e.g., amenity grassland), into pollinator-friendly flower patches. There has also been an increased interest in installing nesting habitat for cavity-nesting insects in the form of artificial traps nests, i.e., ‘bee hotels’ (MacIvor and Packer 2015; MacIvor, 2017). Though primarily installed to recruit cavity-nesting bees, these trap nests also recruit cavity-nesting wasps. These popular interventions are installed to mitigate the negative effects of urbanisation on pollinator communities; however, it is still unclear whether these conservation efforts can help to support urban cavity-nesting bee and wasp communities.

Although cavity-nesting bees and wasps exhibit similar nesting habits both utilise broadly similar floral resources, there is substantial variation in their life histories which may result in differential responses to urbanisation (Wenzel et al., 2020). Cavity-nesting bees rely on floral resources thorough their life cycle by provisioning their young with pollen and feeding on nectar and pollen as adults (Frankie and Thorp, 2009). Dense urban areas may have reduced floral resource diversity (Ellis et al., 2023) and limit the foraging range of bees, which may result in population declines for cavity-nesting bees (Hernandez et al., 2006). Conversely, urban areas have been shown to increase cavity-nesting opportunities in built structures (Banaszak-Cibicka and Zmihorski, 2012; Hinners et al., 2012), which may compensate for habitat loss for cavity nesters, relative to ground nesting bees. (Xie et al., 2022). Cavity-nesting wasps are predatory (e.g., hunting spiders and aphids), parasitoid (e.g., laying eggs inside or on other insects), and cleptoparasitic (i.e., stealing prey or provisions from other Hymenoptera) (Brock et al., 2021) and could therefore suffer more than bees from the negative effects of urbanisation due to these diverse niches (Dürrbaum et al., 2023). For example, parasitoid wasps are more impacted by land cover alterations than their herbivore hosts (Nelson and Forbes, 2014) and Lepidoptera- and spider-predating wasps have a strong negative response to increasing areas of impervious surface (Rocha and Fellowes, 2018). As both these insect communities seem to be at risk in urban ecosystems these is a need to identify conservation interventions that support and enhance both these insect communities by buffering the negative effects of urbanisation. A practical conservation strategy could involve targeted supplementation of their shared resources, such as planting floral patches and installing nesting habitats, but requires demonstration of their effectiveness.

Here, we assess the effect supplementing floral resources has on the uptake of trap nests by cavity-nesting bees and wasps in urban horticultural greenspaces (allotments) along an urbanisation gradient. We experimentally manipulated floral resources by adding nectar-rich flower patches and monitored insect diversity and abundance of bees and wasps in the trap nests over two years. We then compared the colonisation of bees and wasps in trap nests in sites with and without additional floral resources. Within this experimental framework, we asked four questions: 1) Does the provisioning of floral resources enhance the uptake of artificial trap nests? 2) Does urbanisation affect the uptake of artificial trap nests by bees and wasps? 3) Are the effects of urbanisation mitigated by the addition of floral resources? 4) Do bees and wasps differ in their responses to nests, floral additions, and/or urbanisation?

We hypothesise that trap nests uptake will be higher in sites with added floral resources due to the increased availability of nectar for both insect communities. We also predict that urbanisation will negatively affect trap nest uptake due to the associated reduction of resources with increasing urban fabric at a landscape level (Wenzel et al., 2020; Ellis et al., 2023; Liang et al., 2023), but the addition of floral resources should buffer against the severity of this urbanisation gradient. Finally, we hypothesise that there will be variation in responses based on functional group, for example that anthophilous insects (cavity nesting bees) will exhibit stronger responses to the addition of floral resources due to their need to provision their nests with pollen from local flowers. Furthermore, we believe that functional groups with specific non-floral resource needs (cavity nesting wasps) will be more sensitive to urbanisation than cavity nesting bees (Wenzel et al. 2020; Xie et al., 2022; Dürrbaum et al., 2023).

## Methods

### Study system

This study was conducted in urban allotments. Allotments are urban green spaces composed of plots of land rented by an individual or household for growing fruits and vegetables. They have been identified as pollinator and plant hotspots in urban areas (Baldock et al., 2019), containing high insect diversity due to their high floral resources and habitat heterogeneity.

### Experimental design

This study was carried out in 16 allotment sites throughout the 2020 and 2021 growing seasons (March-October) in Leeds, England (53°47’47.33”N, 1°32’52.26”W). The sites were chosen in eight pairs of two geographically clustered sites located along an urbanisation gradient, radiating out from the city centre (Figure 1A). Within each pair there was one site assigned nesting provisioning treatment where trap nests were added to supplement cavity insect nesting resources. The second site was assigned a nesting and a floral provisioning treatment where flower patches (∼100 m^2^) were sown with a nectar-rich seed mix (species list Supplementary Table S1; detailed methods Supplementary Text 1) in March of each year as well as the addition of trap nests (Figure 1).

**Figure 1.**
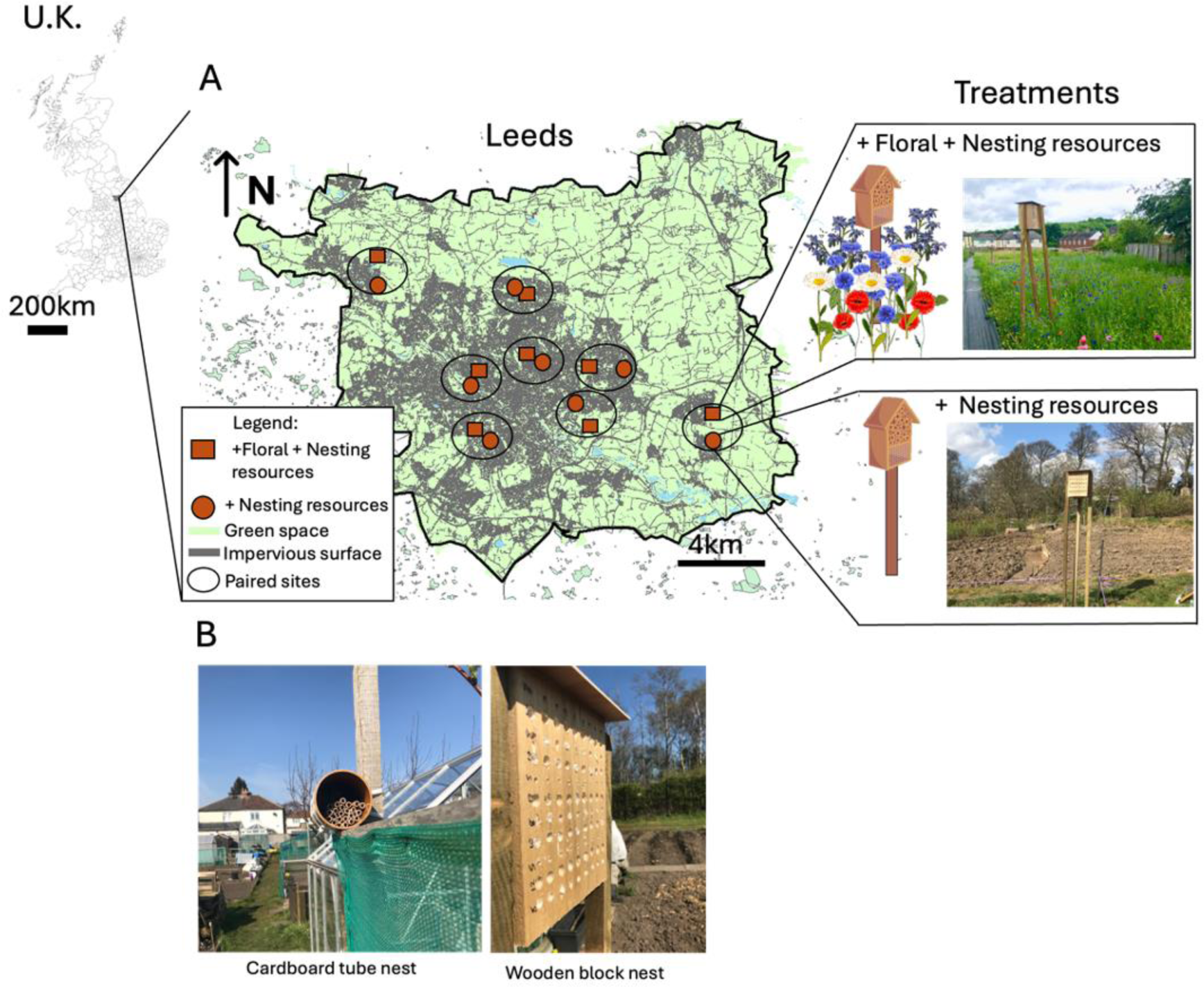
A) left: the city of Leeds location within the U.K., centre: a map of Leeds with allotments sites used in this experiment, showing the configuration of the paired site set-up along an urbanisation gradient (decreasing area of impervious surfaces). Each site received a treatment (+floral + nesting or +floral), which is depicted by the shape of each site location. Right shows the different treatments of our site pairs where floral and nesting resources were added, and the other sites had only nesting resources installed. B) Trap nests type 1: cardboard tube nest, made of 8mm cardboard tubing. Trap nest type 2: wooden block nests were made by drilling 5mm, 8mm,10mm,12mm holes in wooden blocks.

### Trap nests

Trap nest design can influence bee uptake (MacIvor, 2017) and thus we deployed two complementary nest types. We constructed 96 trap nests from PVC pipe filled with cardboard tubes (www.masonbees.co.uk, diameter 8 mm; length 13cm; n= 25 per PVC pipe; FigureFigure 1B), henceforth referred to as cardboard nests. We also constructed wood nests from unprocessed pine wood (25 x 20 x 20 cm) which were drilled with 90 holes of different diameters (equal numbers of 5 mm,8 mm,10 mm,12 mm holes) and 13 cm in length (henceforth referred to as wooden nests; FigureFigure 1B). Each hole was lined with parchment paper before being put in the field so that the larvae could be extracted in winter.

We placed 112 trap nests out for two seasons (2020 and 2021) with nests placed in March and collected in in October (to prevent mould and bacterial colonisation with empty tubes cleaned in 2020 for placement in 2021): all 1.2 m-1.5 m off the ground using wooden stakes, or attaching the nests to existing fences or trees, positioned south-facing in full sun.

In 2020 six cardboard nests and one wooden nest were placed randomly in each of the 16 sites. In 2021 two more wooden nests were added (wooden nests n = 3 per site). All colonised tubes from 2020 and 2021 were overwintered at 4°C and then incubated at 21°C starting early March until all insects had emerged. Bees were identified to species level. Wasps were identified to the lowest taxonomic level possible, resulting in family, genus and morphospecies identification. We quantified three measures of nest uptake: (1) the total number of bees and wasps emerged (abundance), (2) the total number of species of bees and wasps (species richness), (3) functional diversity, depending on their feeding behaviours: pollen feeding (bees), parasitic, cleptoparasitic or predator.

### Landscape mapping

For each allotment site we quantified urbanisation as the area of impervious surface surrounding the allotment sites, which was extracted from UK Ordnance Survey MasterMap© (https://digimap.edina.ac.uk/). Impervious surface included all land-types described as ‘manmade’, including buildings and roads were selected. Circular buffers surrounding the allotment site, with the centre of the circle being the centre of the site, were drawn with a radius of 250 m. The area of impervious surfaces was then exported and used for subsequent analysis.

### Data analysis

All analysis was done in R version 4.12 (R Core Team 2022).

We analysed how insect abundance, species richness and functional diversity were affected by 1) the treatment (nest additions vs nest+floral additions), 2) urbanisation, and 3) the interaction of these terms.

Generalised linear mixed models were constructed using MCMCglmm::MCMCglmm (Hadfield, 2010) as the data were over-dispersed, zero-inflated, and Poisson distribution assumptions were the best match. P-values were derived from the posteriors with standard parameter-expanded priors. We included the season (year 2020 and 2021), nest-type (cardboard nest and wooden nest) and insect taxa (bee and wasps) as model terms, outlined below. Our models followed the same structure:

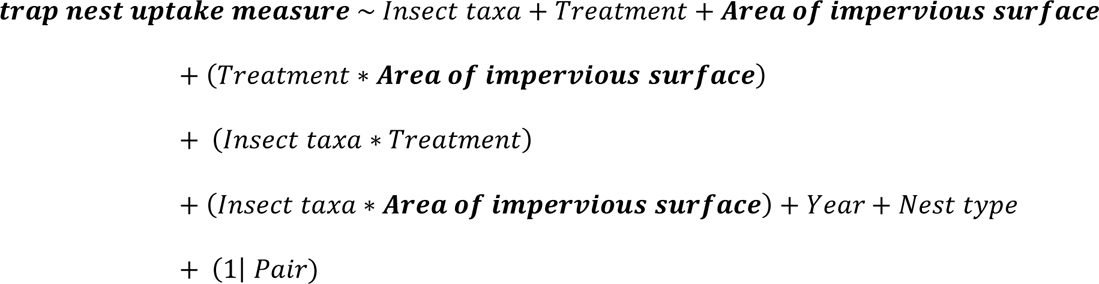

Due to the skewed species distributions, we also separately analysed the four most abundant species (collectively accounting for 74% of the total abundance), pooled for each site, and tested how their abundance varied by urbanisation and our treatment using the same modelling as above, but here we ran four models (with each model having species specific abundance pooled across years as the response variable).

In all analyses with multiple terms and interactions, any non-significant interactions were removed from the final model

## Results

Across the two years a total of 2527 insect emerged from the trap nests (Figure 2). Twenty-five species/species groups of insects were found to colonise the trap nests, including eight species of bees and seventeen types of wasps (Supplementary Table S2). Bees accounted for 39% (n = 979) of the total community, and wasps 59% (n = 1491). The remaining 2% (n = 57) were flies (Diptera) which were subsequently removed from analysis due to low occurrence. The wasp community comprised a range of parasitoids, cleptoparasites and predators. Small parasitic chalcid wasps were the most abundant with 1164 individuals (46% of all insects). The leaf cutter bee (*Megachile ligniseca*) was the second most abundant insect (n = 296) followed by mason bee *Osmia leaiana* (n= 211) and yellow faced bees *Hylaeus hyalinatus* (n = 199) and *Hylaeus communis* (n = 130). The most used cavity size was 8 mm (n = 1191 insects), followed by 5 mm (n = 376), 10mm (n = 140) and 12 mm (n = 13).

**Figure 2.**
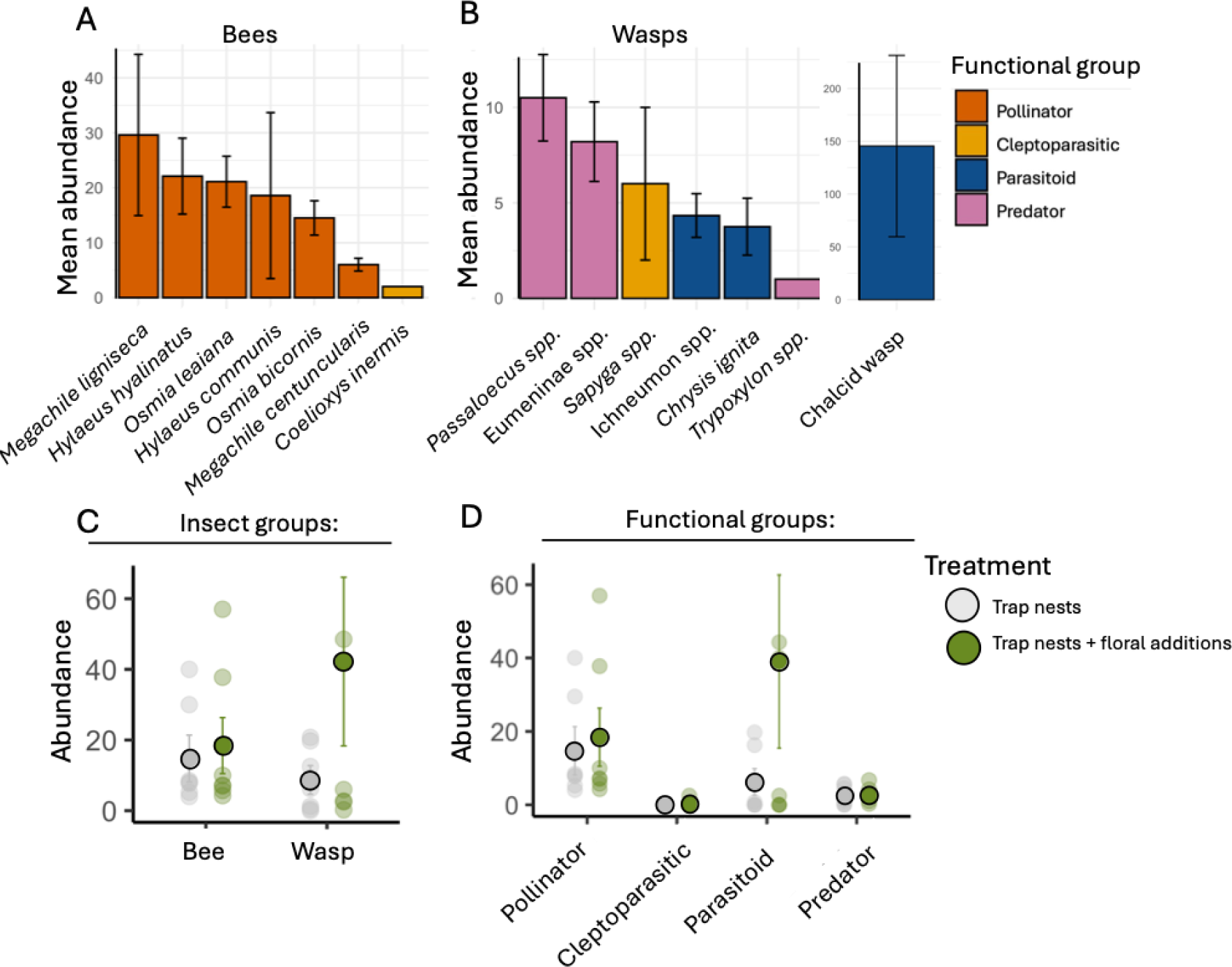
Mean ± SE abundance of cavity-nesting A) bee species and B) wasp assignments across all 16 study allotment sites. Bars are coloured by their assigned functional group. C) This panel shows non-significant (p>0.05) differences in mean (circled dots) insect abundance of bees and wasps across our two treatments. D) Showing non-significant differences in mean insect abundance of functional groups between our two treatments. Error bars show standard error, and background data points are site specific abundances and colours are by treatment: grey = control (+trap nests) and green = supplementation of floral resources (+trap nests + floral).

Uptake was low in the first year (2020) with only 15 of the 112 trap nests (13%) being used but increased to 41 of the 126 trap nests (32%) in 2021. Both bees and wasps showed preference for wooden trap nests compared to cardboard nests with higher species richness (95% CI: 1.91-0.81, p < 0.0001) and abundance (95% CI = 1.82-5.56, p = 0.02) in wooden blocks. There were no differences in nest uptake between our site treatments (floral additions vs control, Figure 2C, D). This was true for abundance (95% CI = 1.93 - 1.70, p = 0.90), species richness (95% CI = −0.52 – 0.47, p = 0.88) of bees and wasps, and across functional groups (Table S3, Figure 2D). With all interactions of insect taxa / functional group x treatment non-significant (p >0.05).

At a landscape level, sites with higher area of impervious surfaces surrounding them had lower uptake of bee nests compared to sites with lower impervious surfaces and the addition of floral resource did not mitigate these declines (Figure 3). Increasing urbanisation had negative effects on overall abundance of bees and wasps (Figure 3A, 95% CI = −2.68--0.13, p = 0.036), with no significant interaction. There was no effect of urbanisation on species richness of bees or wasps (Figure 3B, 95% CI = −1.38-0.15, p = 0.11), or functional groups of bees and wasps (95% CI = −4.51 - 0.26, p = 0.08).

**Figure 3.**
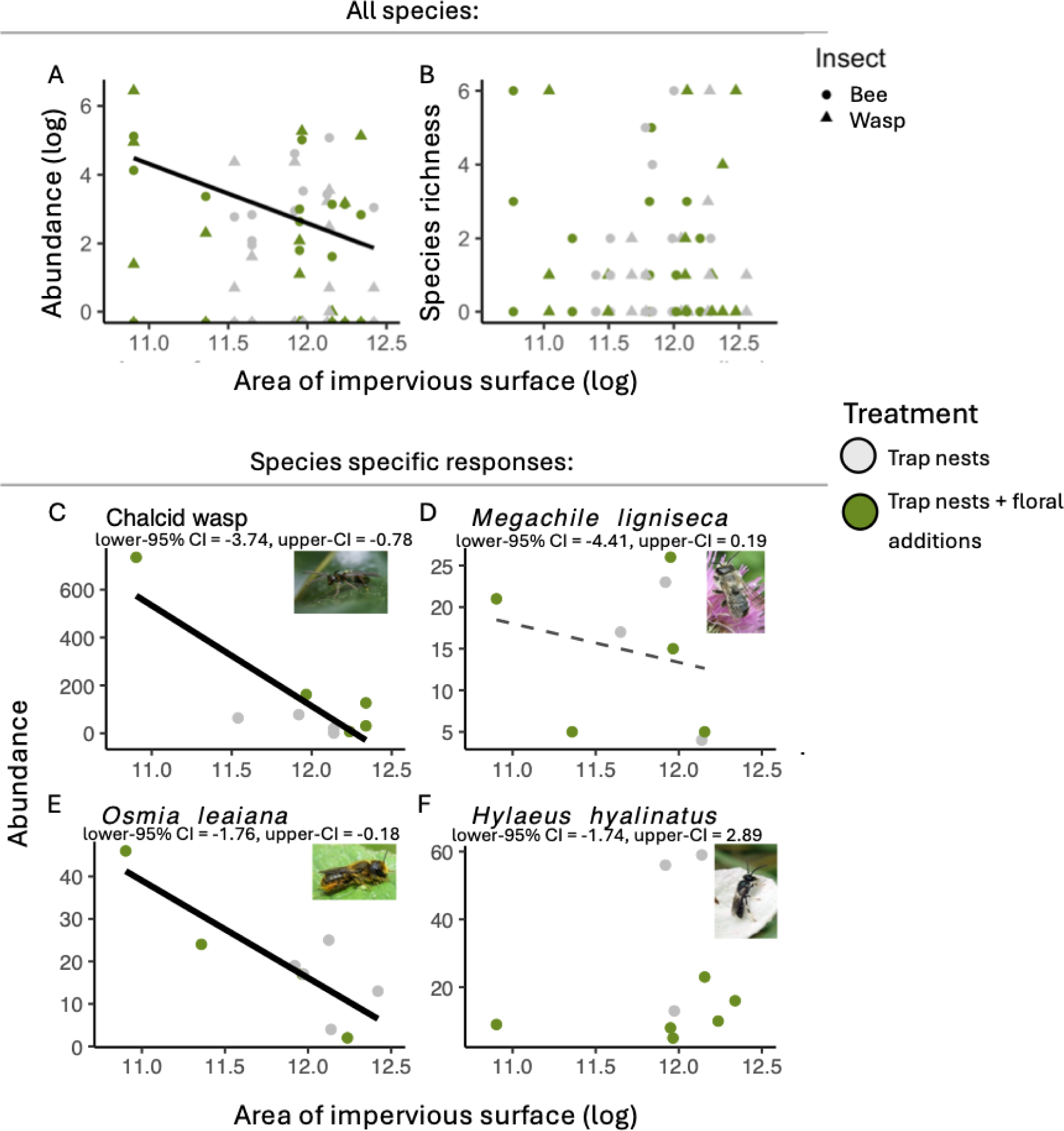
The effect of increasing urbanisation on (A) abundance and (B) species richness of bees (circles) and wasps (triangles) in our treatment sites (nesting and nesting+floral). The effect of increasing urbanisation on abundance of C) Chalcid wasp, D) *Megachile ligniseca*, E) *Osmia leaiana*, F) *Hylaeus hyalinatus*. Solid lines illustrate the significant overall effect urbanisation (no interaction); dashed lines show some evidence of an urbanisation effect (p = 0.08; Supplementary Material Table S4) and no line shows non-significant effect of urbanisation (p >0.08). Photo Credit: Steven Falk.

There was some evidence of species-specific variation in the response of nesting insects to urbanisation. Specifically, the abundance of mason bees (*Osmia leaniana*) and chalcid wasps declined with increasing urbanisation, with no treatment effect (Figure 3, Supplementary Material Table S4). There was also weaker (non-significant) evidence that the leaf cutter bee (*Megachile ligniseca*) abundance was negatively influenced by urbanisation (Figure 3D, Table S4). Yellow-faced bees (*Hylaeus hyalinatus*) showed no response to increasing area of urbanisation (Figure 3F, Supplementary Material Table S4).

## Discussion

A key pollinator conservation management strategy is to improve the provision of floral resources through habitat enhancement (Potts et al., 2005; Spivak et al., 2011). However, the effects of these management tactics are rarely assessed for less charismatic taxa, or in the context of the complex landscape-scale processes associated with urbanisation. Our experimental manipulation of floral and nesting resources along an urbanisation gradient revealed no detectable change in trap nest uptake in sites where floral resources were enhanced. We also found that even with the additional of floral resources, there remained an overall negative effect of surrounding impervious surface area on both wasp and bee colonisation of trap nests. Our results suggest that “pollinator-friendly” floral supplementation may not carry the assumed benefits for cavity-nesting Hymenoptera.

Despite the popularity of planting flower patches in cities, there are surprisingly few studies which have experimentally manipulated floral resources and examined the effects in an urban context. Some studies suggest that wild bees, hoverflies, and honeybees can benefit from the addition of flower patches (Blaauw and Isaacs, 2014) but also have reported no detectable responses (Matteson and Langellotto, 2011). The lack of positive results we report here for cavity-nesting insects underscores the uncertainty surrounding the efficacy of these interventions and their taxon-specific effects. Our results suggest that there may be limitations in the utility of this management intervention potentially driven by both the limitation of pollen and prey resources offered by nectar rich mixes (Potts et al. 2005; Turo and Gardiner, 2019). For example, DNA metabarcoding has demonstrated that pollen from trees is an important resource for cavity-nesting bees (MacIvor et al., 2014; Fernandes et al., 2022, Dürrbaum et al., 2023), and therefore the commonly used flower seed mixes may not provide the right type of resources. For cavity-nesting wasps, the abundance of prey species is necessary to sustain their communities to provision their young (Tscharntke et al., 2005), smaller flower patch additions may be limited in these prey resources.

Prior research has shown that tangible benefits of experimentally adding floral resources are highly context dependent, varying by taxa examined, the scale at which (Matteson and Langellotto, 2011; Blaauw and Isaacs, 2014; Simao et al., 2018) and where the floral additions are made (Wolfin et al. 2023). In our study, we found no discernible benefits from supplementing floral patches on the uptake of trap nests, suggesting that local-scale floral patches may not have the diversity of resources needed to affect the establishment of cavity-nesting insect communities. Our results could also be attributed to the fact that allotments have some of the highest floral diversity and pollinator insect activity in urban areas (Baldock et al., 2019). The high floral density of allotments may have obscured any associated benefits of our floral additions to insect communities due to floral saturation effects (Simao et al., 2018). Had this experiment been conducted in a system where the addition of flower patches significantly boosted overall nectar and pollen production (such as replacing amenity grassland), it’s plausible that there would have been a more pronounced impact on insect visitation (Wolfin et al. 2023).

Cavity-nesting bees and wasps require divergent resources, but contrary to our predictions they did not differ in their sensitivity to urbanisation. This implies that whilst the mechanism driving the decline of these two distinct groups might vary, the effect on nest recruitment was the same (aligning with Holzschuh et al., 2010, Xie et al., 2022). The reduction in cavity-nesting bees is likely due to the associated reducing pollen diversity along an urbanisation gradient (Ellis et al., 2023), especially flowering trees (Nock et al., 2013; Lüttge and Buckeridge, 2020). Our results, therefore, do not support the emerging hypothesis that cavity-nesting bees can persist in highly urban areas due to the nesting opportunities associated with human-made surfaces (Banaszak-Cibicka and Zmihorski 2012; Cane, 2005; Hinners et al., 2012). Our results also contradict Dürrbaum et al. (2023), who show that nest colonisation of cavity-nesting wasps was more strongly negatively affected to urbanisation, compared to cavity-nesting bees. There is a need for more research to pinpoint the mechanisms driving these contrasting results however, the complexity and diversity of cavity-nesting wasp life cycles make it hard to disentangle the direct and indirect drivers of their decline. We believe that measuring the changes in non-floral food sources could shed light on specific functional group limitations. For example, insect predators may be limited due to the reports of decreases in larvae of Lepidoptera in highly urban areas (Boyes et al. 2021). For wasps in non-urban contexts, it has been shown that habitat connectivity/habitat corridors are the best predictors of their community colonisation of trap nests (Bianchi et al., 2006; Holzschuh et al., 2009). Therefore, the associated decreases of habitat connectivity, and increased fragmentation with increasing urbanisation (Liu et al., 2016) is likely driving the declines we see in wasps. Landscape analysis, dispersal and patch-level modelling could help elucidate the roles of patch size and connectivity in urban insect declines, and integrating these findings could then help guide management strategies on the locations, types of intervention and the size of insect-beneficial management strategies.

Considering the ecological value of the ecosystem services provided by cavity-nesting bees and wasp in urban systems (Fernandes et al., 2022; Brock et al., 2021), our report of a 6-fold decline in cavity-nesting insect abundance as urbanisation increases may pose a threat to the functioning of our city ecosystems. While localised conservation efforts have shown promise in mitigating landscape pressures in agricultural ecosystems (Banks-Leite et al., 2020), our research underscores the limitations of small-scale interventions in addressing the complexities of urban systems. Moreover, the lack of knowledge surrounding urban cavity-nesting wasps exacerbates the issue, as their declines likely have far-reaching and poorly understood ecological consequences. Furthermore, it is not yet understood how floral and nesting resource availability shapes parasite and pathogen dynamics in bees and wasps facing urbanisation (Nelson and Forbes, 2014; Cohen et al. 2023) and whether supplementation of floral resources can alter these multi-trophic interactions. This is especially important for urban horticulture as these areas directly depend on pollinators and predators for optimal crop production.

## Conclusion

Our study highlights the complexities involved in effectively implementing urban insect conservation efforts. While adding floral resources is a common strategy, our results indicate that such interventions may not adequately support cavity-nesting bee and wasp communities or mitigate the adverse effects of urbanisation. Furthermore, our findings challenge the widespread assumption that cavity-nesting insects are more resilient in urban environments. The significant decline in these insects that we observed could have profound implications for urban ecosystems, underscoring the urgent need for further research to develop more effective conservation strategies tailored to urban landscapes.

## Statement of authorship

EEE, SAC and JLE conceived and developed the idea. EEE and SAC set up the field experiment. EEE carried out field and lab work. EEE analysed the data. EEE wrote the first draft of the manuscript, and all authors contributed substantially to subsequent revisions.

## Data accessibility statement

All data and R code to reproduce the analysis will be deposited in Dryad. (link to be confirmed once we have a manuscript number).

## Acknowledgments

We thank A Turner, A Maitland for their assistance in the field, and L Rogers, Leeds City Council and allotment secretaries and all the allotment plot holders for site access. This study was supported by a Grantham Centre for Sustainable Futures PhD Studentship, an Engineering and Physical Sciences Research Council (EPSRC) grant to JLE (EP/N030095/1) and a Natural Environment Research Council grant to SAC (NE/R016372/1).

