## Supplementary Information for "Supplementing floral resources does not mitigate the negative effects of urbanisation on cavity-nesting bees and wasps"

Table S3: Summary from generalised linear mixed effect models (MCMCglmm) for the analysis of the effect urbanisation (area of impervious surface surrounding an allotment at a 250m buffer) on the 1) abundance of bees and wasps, 2) abundance of the different functional groups and 3) species richness of bees and wasps found to uptake two types of trap nests (wooden blocks and cardboard) during two years (2020, 2021) and testing if floral additions (Treatment) had main or interactive effects. Significant terms are in bold. ..........4

### Supplementary Tables:

Table S1: Species list of seed mix of EuroFlor and Rigby Taylor Native pollinator and Banquet seed mix (for a more complete list directly contact authors).

| **Species** |
|---|
| *Lotus corniculatus* |
| *Trifolium pratense* |
| *Centaurea cyanus* |
| *Vicia sativa* |
| *Daucus carota* |
| *Papaver rhoeas* |
| *Achillea millefolium* |
| *Myosotis alpestris* |
| *Digitalis purpurea* |
| *Allium schoenoprasum* |
| *Aquilegia vulgaris* |
| *Borago officinalis* |
| *Calendula officinalis* |

Table S2: List of cavity nesting bees and wasps, their species or morphotypes, feeding behaviours that colonised artificial trap nests during 2020 and 2021 in sixteen allotment sites in Leeds.

| Year | Insect | Morphotype | Species | Feeding behaviour | Abundance |
|---|---|---|---|---|---|
| 2020 | | | | | |
| | bee | | *Osmia bicornis* | **Pollinator** | 42 |
| | bee | | *Osmia leaiana* | **Pollinator** | 19 |
| | wasp | Mason wasp 1 | *Ancistrocerus* sp. | Lepidoptera caterpillars | 31 |
| | wasp | Resin sealing wasp 1 | *Passaloecus/Pemphredon* | Spider/ aphid hunter | 3 |
| | wasp | Ruby tailed wasp | *Chrysis ignita* agg. | Kleptoparasites | 1 |
| 2021 | | | | | |
| | bee | | *Megachile lignisea* | **Pollinator** | 296 |
| | bee | | *Hylaeus hyalinatus* | **Pollinator** | 199 |
| | bee | | *Osmia leaiana* | **Pollinator** | 148 |
| | bee | | *Hylaeus communis* | **Pollinator** | 130 |
| | bee | | *Osmia bicornis* | **Pollinator** | 74 |
| | bee | | *Megachile centuncularis* | **Pollinator** | 20 |
| | bee | | *Coelioxys inermis* | **Pollinator** | 2 |
| | wasp | Chalcid wasp 1 | *Monodontomerus* sp. | Bee parasitoid | 1098 |
| | wasp | Chalcid wasp 2 | *Monodontomerus* sp. | Bee parasitoid | 128 |
| | wasp | Resin sealing wasp 1 | *Passaloecus/Pemphredon* | Spider/ aphid hunter | 101 |
| | wasp | *Gasteruption jaculator* | Gasteruption jaculator | Bee parasitoid | 21 |
| | wasp | Ruby tailed wasp | *Chrysis ignita* agg. | Kleptoparasites | 14 |
| | wasp | Ichneumon 1 | | Bee parasitoid | 10 |
| | wasp | Mason wasp 1 | *Ancistrocerus* sp. | Lepidoptera | 8 |
| | wasp | Ichneumon 2 | | Bee parasitoid | 2 |
| | wasp | Ichneumon 3 | | Bee parasitoid | 2 |
| | wasp | Mason wasp 2 | *Ancistrocerus* sp. | Lepidoptera caterpillars | 1 |
| | wasp | Mason wasp 3 | *Ancistrocerus* sp. | Lepidoptera caterpillars | 1 |
| | wasp | Ichneumon 4 | | Bee parasitoid | 1 |
| | wasp | Ichneumon 5 | | Bee parasitoid | 1 |
| | wasp | Ichneumon 6 | | Bee parasitoid | 1 |
| | wasp | Ichneumon 7 | | Bee parasitoid | 1 |
| | wasp | Resin sealing wasp 2 | *Passaloecus/Pemphredon* | Spider/ aphid hunter | 1 |

Table S3: Summary from generalised linear mixed effect models (MCMCglmm) for the analysis of the effect urbanisation (area of impervious surface surrounding an allotment at a 250m buffer) on the 1) abundance of bees and wasps, 2) abundance of the different functional groups and 3) species richness of bees and wasps found to uptake two types of trap nests (wooden blocks and cardboard) during two years (2020, 2021) and testing if floral additions (Treatment) had main or interactive effects. Significant terms are in bold.

| | Post mean | 95% CI | u-95%CI | effective sample | p-value |
|---|---|---|---|---|---|
| Abundance (bees vs wasps) | | | | | |
| (Intercept) | 29.32665 | -0.08964 | 61.48786 | 8040 | 0.0546 |
| Year (2021) | 4.17863 | 2.32072 | 6.14854 | 3979 | **<1e-04** |
| log(area of impervious surface) | -2.67337 | -5.28008 | -0.12577 | 7375 | **0.0373** |
| Nest_type (Cardboard) | -2.44465 | -4.28098 | -0.65631 | 6650 | **0.0068** |
| Insect taxa (wasp) | -0.38295 | -2.14278 | 1.35100 | 8477 | **0.6594** |
| Treatment (+Floral +Nesting) | -0.12146 | -1.90760 | 1.67663 | 8567 | 0.8903 |
| Abundance (functional groups) | | | | | |
| (Intercept) | 21.77004 | -7.62966 | 49.25606 | 7013 | 0.123505 |
| Year (2021) | 4.38198 | 2.59432 | 6.22323 | 1642 | **<1e-04** |
| log(area of impervious surface) | -2.04388 | -4.49110 | 0.27718 | 6738 | **0.084124** |
| Nest_type (Cardboard) | -3.00751 | -4.73338 | -1.35315 | 2755 | **0.000619** |
| Functional group - Cleptoparasitic | -7.20740 | -10.28465 | -4.07010 | 1608 | **<1e-04** |
| Functional group - Parasitoid | -2.77964 | -4.93172 | -0.71095 | 4145 | **0.004330** |
| Functional_group - Predator | -1.31166 | -3.27660 | 0.46156 | 8546 | 0.163918 |
| Treatment (+Floral +Nesting) | 0.08432 | -1.53628 | 1.64675 | 6595 | 0.910103 |
| Species richness (bees vs wasps) | | | | | |
| (Intercept) | 4.87889 | -4.42689 | 14.10864 | 1614.9 | 0.301 |
| Year (2021) | 2.01512 | 1.37805 | 2.70063 | 623.3 | **<1e-04** |
| log(area of impervious surface) | -0.52179 | -1.32754 | 0.22676 | 1550.6 | 0.174 |
| Nest_type (Cardboard) | -1.40353 | -1.97808 | -0.80953 | 527.6 | **<1e-04** |
| Insect taxa (wasp) | -0.04526 | -0.52445 | 0.46833 | 1546.5 | 0.877 |
| Treatment (+Floral +Nesting) | 0.08118 | -0.46484 | 0.59081 | 1321.6 | 0.726 |

Table S4: Summary from generalised linear mixed effect models (MCMCglmm) for the analysis of the effect urbanisation (area of impervious surface surrounding an allotment at a 250m buffer) on the four individual species of bees and wasps found to uptake trap nests during two years (aggregated).

**Species**

*Osmia leaiana*

| | Post mean | 95% CI | u-95%CI | effective sample | p-value |
|---|---|---|---|---|---|
| (Intercept) | 14.1145 | 5.1365 | 23.6204 | 6326 | 0.0113 |
| log(area of impervious surface) | -0.9560 | -1.7903 | -0.1846 | 6229 | **0.0239** |

*Megachile ligniseca*

| | Post mean | 95% CI | u-95%CI | effective sample | p-value |
|---|---|---|---|---|---|
| (Intercept) | 26.93631 | -0.14917 | 53.25422 | 9237 | 0.0524 |
| log(area of impervious surface) | -2.06207 | -4.40117 | 0.08522 | 9233 | 0.0687 |

*Hylaeus hyalinatus*

| | Post mean | 95% CI | u-95%CI | effective sample | p-value |
|---|---|---|---|---|---|
| (Intercept) | -8.0697 | -34.2207 | 17.4351 | 9341 | 0.485 |
| log(area of impervious surface) | 0.8989 | -1.2470 | 3.0549 | 9360 | 0.366 |

Chalcid wasp

| | Post mean | 95% CI | u-95%CI | effective sample | p-value |
|---|---|---|---|---|---|
| (Intercept) | 31.3992 | 14.5091 | 50.0067 | 8597 | **0.0103** |
| logMM_250m | -2.2944 | -3.8987 | -0.9017 | 8624 | **0.0151** |

#### Supplementary Text:

#### **Text S1:** Flower patch addition methodology:

Site preparation (March):

- Cut vegetation down to 5-10cm and turn the soil over in an area of 100$m^2$
- Remove persistent weeds

Seed bed preparation and sowing (April-May):

- Rake over soil and remove debris and stones.
- Sow 3$g/m^2$ of seeds evening throughout the patch (on a calm, dry day).
- Roll over the seeds lightly to maximise germination.
